# Properties of the *Ureaplasma parvum* SMC protein related to its interaction with DNA

**DOI:** 10.1101/2024.04.14.589448

**Authors:** Natalya A. Rumyantseva, Vladimir M. Shutov, Dina G. Belenkaia, Aleksandr A. Alekseev, Natalia E. Morozova, Alexey D. Vedyaykin

## Abstract

SMC (Structural Maintenance of Chromosomes) ATPase proteins are integral components of complexes bearing the same name, crucial for the spatial organization of DNA across diverse life forms, spanning bacteria, archaea, and eukaryotes. It is proposed that in bacteria, SMC complexes facilitate DNA compaction through loop extrusion and aid in the segregation of daughter nucleoids. In this paper the properties of the SMC ATPase protein from *Ureaplasma parvum* were investigated by using a spectrum of methods, including conventional biochemical methods as well as advanced single-molecule techniques. Our findings reveal distinctive properties of this protein compared to its extensively studied homologue from *Bacillus subtilis*. Notably, our results suggest that *U. parvum* SMC ATPase facilitates DNA compaction even in the absence of ATP.

## Introduction

SMC (Structural Maintenance of Chromosomes) protein complexes are ubiquitous across the biological spectrum, from bacteria to eukaryotes, playing pivotal roles in orchestrating the spatial arrangement of DNA [1, 2]. These complexes typically comprise three primary components: a dimer of SMC ATPase proteins, a kleisin, and auxiliary proteins such as KITE or HAWK [3]. SMC ATPase proteins form a dimeric structure — heterodimeric in eukaryotes and homodimeric in prokaryotes — characterized by a V-shape, with each arm of the dimer composed of coiledcoils spanning approximately 50 nm. Notably, SMC proteins exhibit ATPase activity, which is usually modulated upon binding to DNA. Kleisin proteins bind to the arms of the SMC dimer, facilitating the formation of a closed conformation, while auxiliary proteins like KITE and HAWK interact with SMC and kleisin, playing regulatory and auxiliary roles. Central to the functionality of SMC complexes is their capacity for loop extrusion, wherein they actively generate DNA loops, categorizing them as a subclass of DNA translocases [1]. Direct demonstration of loop extrusion has been achieved for eukaryotic SMC complexes — condensins [4], cohesins [5], and Smc5/Smc6 complexes [6] — under *in vitro* conditions at the single-molecule level.

Bacterial genomes harbor diverse SMC complexes, including SMC-ScpAB [7], MukBEF [8], Wadjet [9], and others [10], with SMC-ScpAB of *Bacillus subtilis* and MukBEF of *Escherichia coli* being the most extensively studied. In these complexes, SMC and MukB serve as ATPases, while ScpA and MukF function as kleisins, and ScpB and MukE act as auxiliary proteins of the KITE type [3]. SMC-ScpAB and MukBEF complexes play crucial roles in bacterial DNA segregation, albeit through distinct mechanisms. The SMC-ScpAB complex of *B. subtilis* associates with the bacterial chromosome, facilitated by ParB, and specifically binds to DNA at *parS* sites [11]. Conversely, the MukBEF complex of *E. coli* exhibits nonspecific binding to the chromosome [12]. Moreover, SMC-ScpAB-mediated action results in the formation of a crescent-shaped nucleoid, while the MukBEF complex yields a ring-shaped nucleoid. Notably, while the ability of bacterial SMC complexes to extrude loops under *in vitro* conditions remains unproven, *in vivo* findings [13] and some single-molecule experiments conducted *in vitro* [14, 15] support the presence of such capability.

SMC complexes likely play indispensable roles in DNA segregation in certain bacterial contexts. For instance, the MukBEF complex of *E. coli* is essential during periods of rapid growth [15]. Furthermore, their significance is underscored by their presence in the reduced genomes of Mollicutes; genes encoding proteins of the SMC-ScpAB complex persisted even after directed deletion of approximately half of the genes in a synthetic “minimal bacterium” [16]. Investigating the properties of SMC complexes in Mollicutes is of particular interest, given the potential for simpler organizational schemes compared to bacteria with “complete” genomes, possibly leading to simpler regulatory mechanisms. Additionally, the functions of Mollicutes SMC complexes appear to exhibit less overlap with those of other protein complexes involved in DNA segregation compared to more complex bacteria. This study is devoted to SMC ATPase protein from the SMC-ScpAB complex of *U. parvum*, revealing several notable distinctions between this protein and its homologue in *B. subtilis*.

## Materials and Methods

### Creation of Genetic Construct for *U. parvum* SMC Protein Production

The gene encoding the SMC protein of *U. parvum* was synthesized *de novo* by IDT DNA with codon optimization for *E. coli* expression. Utilizing the Codon Optimization Tool (IDT DNA), the nucleotide sequence of the gene was changed for efficient expression in *E. coli*. The synthesized gene was integrated into the pJET1.2 plasmid and verified by Sanger sequencing. The resulting plasmid was transformed into *E. coli* XL-1 Blue strain. Subsequently, the gene of interest, verified by sequencing, was amplified from the pJET1.2-based plasmid using specific primers with NdeI and XhoI restriction sites added, and subsequently incorporated into the pET21a plasmid via standard restriction and ligation cloning. Primers synthesis was performed by Evrogen, and the resulting pET21a-based plasmids were also subjected to sequencing verification.

### Production and Purification of *U. parvum* SMC Protein

*E. coli* BL21 strain served as the host for *U. parvum* SMC protein production. Confirmation of protein synthesis was achieved by observing the corresponding band on SDS-PAGE gel. Identification of the protein in the band was accomplished through mass spectrometry, employing the peptide fingerprinting method as described in reference [17].

Recombinant SMC protein was expressed in a 1-liter culture. Bacteria were cultivated in a thermoshaker at 37°C until reaching an optical density at 600 nm (OD600) of 1. Gene expression induction was initiated by adding IPTG to a final concentration of 1 mM, followed by overnight incubation at 22°C in a thermoshaker. Cells were harvested by centrifugation, the supernatant was decanted, and cells were resuspended in buffer A (50 mM Tris-HCl, 300 mM NaCl, 10 mM MgCl_2_, and 1 mM PMSF, pH 8.0). Subsequent centrifugation at 3600 g and 4°C for 30 minutes yielded a cell pellet, which was then frozen at -80°C.

The frozen cell pellet was thawed on ice and resuspended in buffer A. Cell disruption was performed on ice using a sonicator, with prior addition of lysozyme (1 mg/mL). The resultant lysate was subjected to centrifugation at 12000 g for 20 minutes at 4°C, and the supernatant was collected and filtered. This lysate was applied onto a 1 mL HisCap Smart 6FF column (Smart-Life Sciences) using an Akta Purifier 10 chromatograph (GE Healthcare). Subsequent washing steps were carried out with mix of buffer A and buffer B (buffer A + 500 mM imidazole), with the imidazole concentration being incrementally increased. Fractions (1 mL) were collected from each peak of the chromatogram, flash-frozen in liquid nitrogen, and stored at -80°C. The obtained samples underwent SDS-PAGE electrophoresis analysis. SMC protein concentration was estimated by absorbance at 280 nm (OD280), with correction for the molecular weight (MW = 112 kDa) and molar extinction coefficient (ε = 88443 cm∧-1*M∧-1) of *U. parvum* SMC protein.

### Electrophoretic Mobility Shift Assay (EMSA) in Polyacrylamide Gel

Two fluorescently labeled DNA molecules, a 110 bp fragment from the human GRIN2B gene and a 110 bp sequence from the *U. parvum* genome (with a GC content of approximately 20%), designated as MTA (the gene encoding the 5’-methylthioadenosine/S-adenosylhomocysteine nucleosidase protein), were utilized to analyze DNA electrophoretic mobility shift in polyacrylamide gel via EMSA. Fluorescently labeled molecules were generated by PCR using primers, one of which was labeled with Cy3 at the 5′-end. Partially single-stranded DNA molecules were obtained by subjecting double-stranded DNA to melting at 95°C followed by rapid cooling on ice.

Reaction mixtures (20 μL) comprised 20 mM Tris-HCl, pH 7.5, 7.5 mM KCl, 2.5 mM MgCl_2_, 1 mM DTT, and varying concentrations of SMC protein (final concentrations ranging from 0.065 to 1.04 μM). After centrifugation at 16000 g for 15 min, the supernatants were withdrawn and incubated at 37°C for 5 min. Subsequently, fluorescently labeled DNA (either partially single-stranded or fully double-stranded molecules) was added to each sample to a final concentration of 5 nM, with DNA-to-protein ratios ranging from 1:10 to 1:200.

Following dark incubation at 37°C for 30 min, glycerol was added to the mixture up to 10%, and samples were subjected to native electrophoresis in a 5% polyacrylamide gel in TBE buffer at 70 V for 50 min on ice. Pre-equilibration of the gel in the same buffer was performed at 80 V for 30 min. Gel visualization was carried out using the ChemiDoc gel documentation system (Bio-Rad) with Cy3 dye channel selection.

### Electrophoretic Mobility Shift Assay (EMSA) in Agarose Gel

DNA substrates utilized were circular or linearized plasmid pUC19 dual (9396 bp) [18]. For most subsequent experiments, the following protocol was adopted. SMC protein was centrifuged at 21130 g for 10 min at 4°C to remove insoluble protein, and the supernatant was collected. Subsequently, 4 μg of SMC protein was added to the reaction mixture based on one of the buffers (as described below), followed by the addition of 200 ng of circular or linear pUC19 dual plasmid (DNA:protein ratio of 1:1000). Two control samples were prepared: one lacking SMC protein but containing the same DNA amount as the other reaction mixtures, and another lacking DNA but containing SMC protein at the same concentration as the tested samples. ATP was added to the samples (if necessary), followed by incubation for 40 minutes at 37°C.

Samples were loaded onto a 0.7% agarose gel in Tris-acetate buffer, and electrophoresis was conducted at a constant voltage of 80 V for 25 minutes. Post-electrophoresis, the gel was incubated in TAE containing ethidium bromide (1/10000) for DNA visualization.

Several buffers with varying compositions were employed for different experiments:

1. To assess the effect of ATP on SMC protein’s DNA-binding properties, identical reaction mixtures based on pH 7 buffer (20 mM Tris-HCl, 1 mM DTT, 7.5 mM KCl, 2.5 mM MgCl_2_, pH 7) were prepared, differing in ATP concentration: 0 mM, 2.5 mM, 5 mM, 7.5 mM, 10 mM.
2. To evaluate the impact of buffer pH on SMC protein’s DNA-binding properties, reaction mixtures based on buffers with different pH values (pH 6, 7, 8, 9, 10) were prepared. For pH 7, 8, 9, 10, buffers comprised 20 mM Tris-HCl, while for pH 6, the buffer was based on 20 mM MES. All mixtures contained 1 mM DTT, 7.5 mM KCl, 2.5 mM MgCl_2_, and 5 mM ATP.
3. To assess the influence of KCl concentration on SMC protein’s DNA-binding properties, mixtures based on pH 6 buffer (20 mM MES, 1 mM DTT, 2.5 mM MgCl_2_, 5 mM ATP, pH 6) were utilized, with varying concentrations of KCl salt: 0 mM, 25 mM, 50 mM, 75 mM, 100 mM.
4. To evaluate the effect of MgCl_2_ concentration on SMC protein’s DNA-binding properties, buffer-based mixtures at pH 6 (20 mM MES, 1 mM DTT, 7.5 mM KCl, 5 mM ATP, pH 6) with different MgCl_2_ concentrations were used: 0 mM, 10 mM, 50 mM, 100 mM, 200 mM.
5. Additionally, the impact of DNA:SMC ratio on SMC-DNA interaction was determined. Reaction mixtures based on pH 6 buffer (20 mM MES, 1 mM DTT, 7.5 mM KCl, 2.5 mM MgCl_2_, 5 mM ATP, pH 6) were prepared, with varying DNA:SMC ratios (1:1000, 1:800, 1:600, 1:400, 1:200), while maintaining a constant DNA amount of 200 ng.

### Measurement of ATPase Activity

The reaction mixture consisted of 20 mM MES (pH 6), 7.5 mM KCl, 2.5 mM MgCl_2_, 1 mM DTT, 656 nM SMC protein (1.47 μg), 1.64 nM DNA (ring plasmid pUC19dual - 200 ng), and 1 mM ATP. The mixture was incubated at 37°C for specific time intervals (5, 10, 15, 20 min). Subsequently, 20 μl of the reaction mixture was transferred into a 96-well flat-bottom plate. 200 µl of color reagent (consisting of 1 part 4.2% ammonium molybdate, 3 parts 0.045% malachite green solution, and 1/50 part 2% Nonidet P-40 solution) was added to the samples and mixed. After 1 min, 20 μl of 34% sodium citrate solution was added and stirred. The absorbance of the solutions at 620 nm wavelength was analyzed using a CLARIOstar platereader spectrophotometer. To determine the concentration of free phosphate in the samples, a calibration line corresponding to a standard with known values of free phosphate concentration was utilized. The standard used was 1 mM NaH_2_PO_4_, which was diluted with water to concentrations of 0, 50, 100, 150, 200 μM.

### Molecular Combing

For molecular combing, λ phage DNA, 48502 bp in length, pre-stained with YOYO-1 fluorescent dye, was employed. The reaction mixture consisted of 20 mM Tris-HCl (pH 7.5), 7.5 mM KCl, 2.5 mM MgCl_2_, 1 mM DTT, 1 mM ATP, 1.3 μM SMC, and 0.3 nM λ DNA. Reaction mixtures devoid of ATP or SMC protein served as controls. Incubation of the reaction mixtures occurred in a thermostat at 37°C for 30 min. Following this, 100 mM sodium acetate buffer (pH 5.5) was added to each mixture to facilitate subsequent DNA application to coverslips via the molecular combing method [19].

Pre-cleaned (via plasma cleaning in the PlasmaCleaner) 22×22 mm coverslips were hydrophobically coated by evenly applying 200 μl of polystyrene solution in benzene using a spin-coater. Subsequently, the coverslips were washed with water. The polystyrene-coated coverslip was incubated in buffer with the reaction mixture for 5 min, after which the glass was slowly pulled through a meniscus of solution to unfold and anchor the DNA molecules on the glass. Visualization and capture of the dye-labeled DNA were accomplished using standard fluorescence microscopy, specifically the YFP fluorescence channel.

### DNA Manipulation by Laser Tweezers

Optical trapping experiments were conducted to analyze SMC protein binding to individual double-stranded DNA molecules. DNA manipulation was performed within a five-channel microfluidic system (uFlux, Lumicks), utilizing two polystyrene microspheres coated with streptavidin (Spherotech, 2.1 μm diameter) held by two optical traps, according to the methodology outlined in references [20, 21]. A 11070 bp long linear DNA molecule (prlC) served as the substrate. The solution comprised Tris-HCl buffer (pH 7.5, 25 mM), 7.5 mM KCl, 2.5 mM MgCl_2_, and 0.02% BSA.

## Results and Discussion

### SMC protein from *U. parvum* is capable of interacting with double-stranded DNA

The analysis of electrophoretic mobility shift assays revealed that *U. parvum* SMC forms complexes with double-stranded DNA. Both polyacrylamide and agarose gel experiments demonstrated the ability of *U. parvum* SMC protein to bind to DNA (Figure 1). Notably, the protein exhibited a preference for double-stranded DNA, as the presence of single-stranded sites on the DNA molecule, generated by rapid annealing of the PCR product, did not enhance the interaction with SMC. Similarly, any detectable shift was absent in the case of single-stranded M13 phage DNA analyzed in agarose gel (data not shown). This finding contrasts with previous observations in a similar study [22], where the homologous SMC protein from *B. subtilis* was reported to bind preferentially to single-stranded DNA, exhibiting minimal affinity for double-stranded DNA.

**Figure 1:**
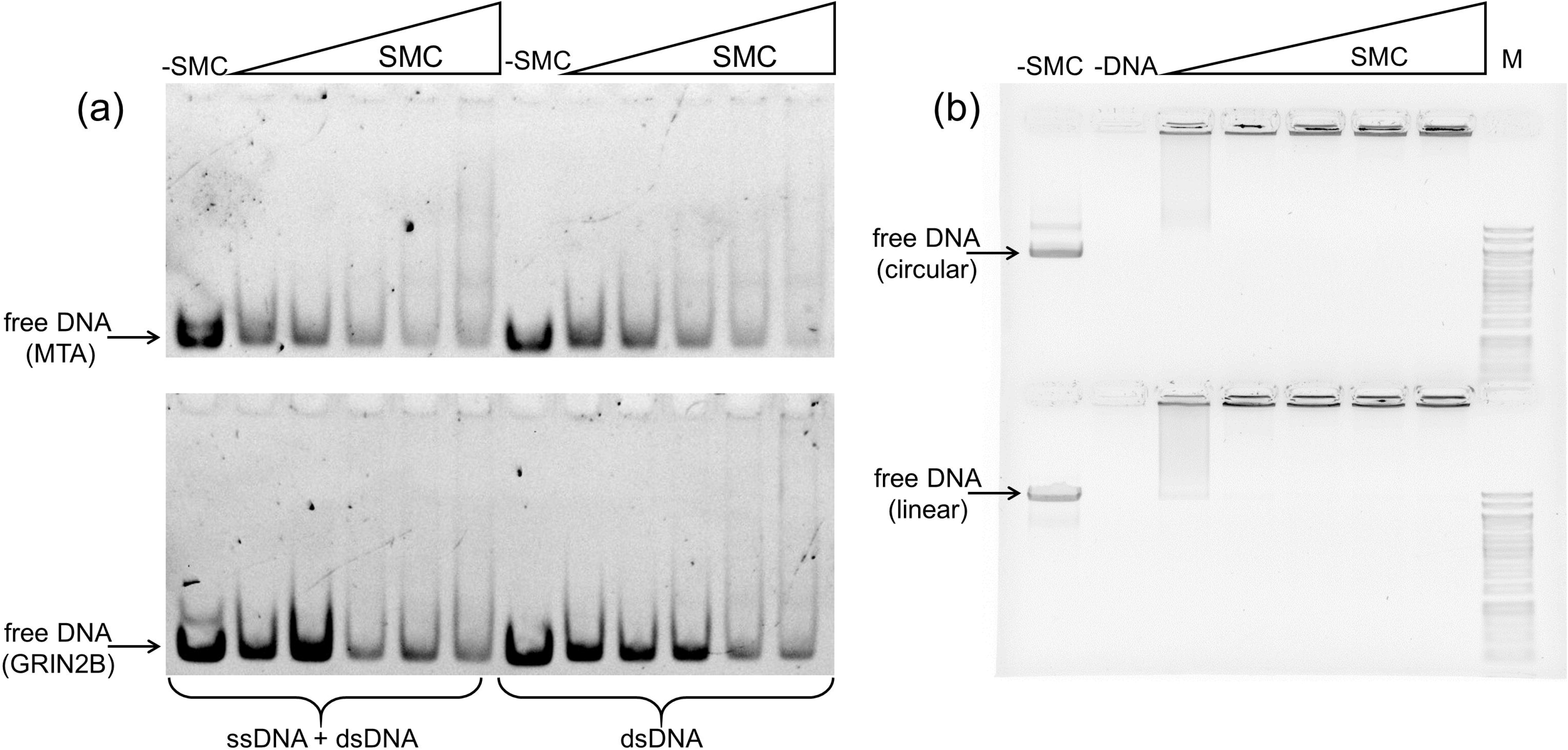
The effect of *U. parvum* SMC on the electrophoretic mobility of DNA. (a) Binding of *U. parvum* SMC protein to linear DNA fragments visualized in a polyacrylamide gel. The binding of SMC to a mixture of single- and double-stranded DNA obtained by rapid annealing of the PCR product (ssDNA + dsDNA) and to double-stranded DNA (dsDNA) is shown. Top: Binding of *U. parvum* SMC protein to a 110 bp MTA fragment of the *U. parvum* genome. Bottom: Binding of *U. parvum* SMC protein to a 110 bp fragment of the human GRIN2B gene. Samples with a stepwise increase in the amount of SMC from 0 (-SMC) to the maximum amount according to the ratios of DNA:SMC molecules (1:10, 1:20, 1:50, 1:100, 1:200, respectively) are shown. (b) Binding of *U. parvum* SMC protein to the circular and linear form of DNA (9396 bp plasmid pUC19-dual) visualized in an agarose gel. Top: Binding of SMC protein to the circular form of the plasmid. Bottom: Binding of the SMC protein to the linear form of the plasmid. Samples with a stepwise increase in SMC amount from 0 (-SMC) to the maximum amount according to the ratios of DNA:SMC molecules (1:200, 1:400, 1:600, 1:800, 1:1000, respectively) are shown. M – marker (Thermo Scientific GeneRuler 1kb DNA Ladder), -DNA – control sample without DNA addition. -SMC – control sample without SMC addition

In *B. subtilis*, the loading of SMC onto the chromosome by ParB at *parS* sites ensures specific binding of SMC to the origin of replication [11]. Conversely, the SMC-like MukBEF complex in *E. coli* exhibits strong, albeit nonspecific, binding to double-stranded DNA [23]. Our observation that *U. parvum* SMC interacts with double-stranded DNA without additional factors suggests a potential nonspecific binding mechanism. Indeed, the results support the notion of nonspecific DNA binding by *U. parvum* SMC, as evidenced by the continuous smear-like pattern observed on the gel tracks, along with complexes entrapped in the wells, likely due to the formation of large SMC-DNA complexes. Notably, the binding of SMC to DNA in our experiments occurred in the absence of ATP.

Surprisingly, *U. parvum* SMC protein demonstrated comparable interactions with both circular and linear forms of DNA (Figure 1b). This result contrasts with observations for other SMC proteins, which typically exhibit a preference for circular DNA. Various SMC complexes, such as condensins, cohesins, and Smc5-Smc6, have been reported to display topological binding to DNA, with a stronger affinity for circular DNA compared to linear DNA [24-26]. For instance, RecN, an SMC-like protein, exhibits a significantly stronger interaction with circular DNA, especially single-stranded DNA, compared to linear DNA under similar experimental conditions [27]. The lack of published results on the properties of *B. subtilis* SMC protein under conditions similar to our experiments precludes a direct comparison.

The identical binding of *U. parvum* SMC protein to circular and linear DNA suggests the formation of unusually stable SMC/DNA complexes that do not dissociate under the experimental conditions employed. The high stability of such complexes may be attributed, in part, to the relatively low ATPase activity of *U. parvum* SMC protein (see below). Further investigations are warranted to elucidate the underlying mechanisms governing the binding of *U. parvum* SMC to DNA and its preference for specific DNA topologies.

### Enhancement of DNA Binding by ATP in *U. parvum* SMC

The interaction of *U. parvum* SMC with DNA is enhanced by the addition of ATP. Analysis of the shift in electrophoretic mobility of different forms of double-stranded DNA (circular supercoiled and linear) in the presence of SMC protein in an agarose gel showed similar results: SMC interacts with DNA and forms complexes with it even in the absence of ATP in the solution. It was possible to demonstrate the dependence of the DNA-binding properties of the investigated protein on the ATP concentration (see Figure 2a). Increasing the ATP concentration enhanced the interaction of SMC protein with DNA: more active formation of protein/DNA complexes in the gel well was observed with increasing ATP concentration (see Figure 2a). At high ATP concentrations (more than 5 mM), most of the 9396 bp molecules of plasmid pUC19 dual appeared to be immobilized in lanes. Since at ATP concentration of 5 mM most of the DNA was complexed in SMC, this concentration was used in the following experiments.

**Figure 2:**
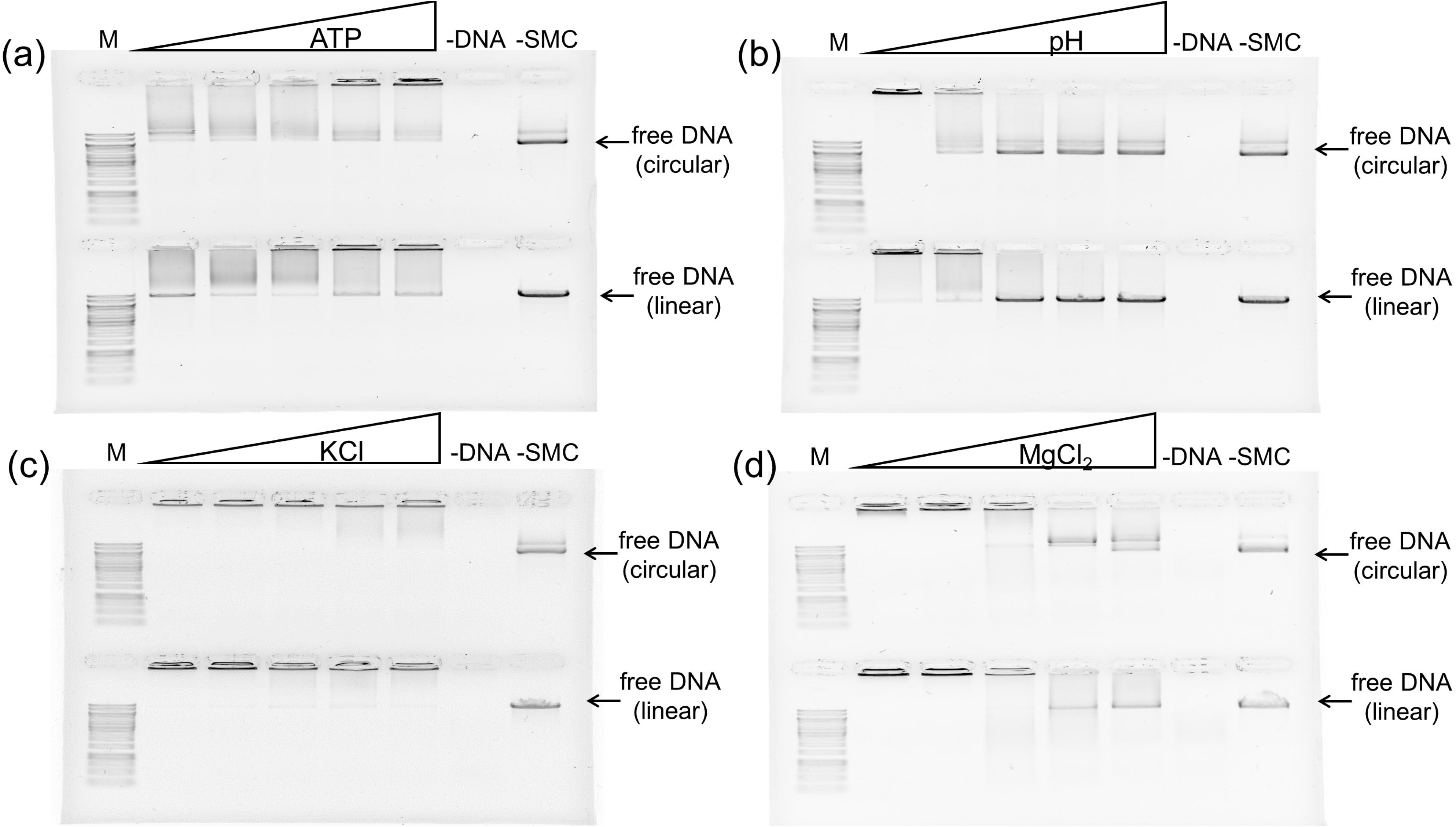
Impact of buffer composition on the interaction of *U. parvum* SMC protein with DNA (9396 bp plasmid pUC19-dual) visualized in agarose gel via EMSA. (a) Effect of ATP concentration (0, 2.5, 5, 7.5, and 10 mM). (b) Effect of solution pH (pH 6, 7, 8, 9, and 10). (c) Effect of KCl concentration (0, 25, 50, 75, and 100 mM). (d) Effect of MgCl_2_ concentration (0, 10, 50, 100, and 200 mM). The top part of each part depicts SMC protein binding to circular plasmid, while the bottom part shows binding to linear plasmid. M: Marker (Thermo Scientific GeneRuler 1kb DNA Ladder), -DNA: Control sample without DNA, -SMC: Control sample without SMC

### Effect of pH, KCl, and MgCl_2_ Concentrations on DNA Binding

The interaction of *U. parvum* SMC with DNA was weakened with increasing pH and concentration of KCl, MgCl_2_. Increasing pH above 7 weakened the interaction of SMC protein with DNA: less active formation of SMC/DNA complexes in the gel well was observed at higher pH (see Figure 2b). Since pH = 6 was found to be optimal for protein-DNA binding, it was the pH = 6 buffer that was used in subsequent experiments. It was found that increasing the concentration of KCl starting at 50 mM inhibited the interaction between DNA and SMC (see Figure 2c). It was also found that increasing the concentration of MgCl_2_ starting at 50 mM inhibited the interaction between DNA and SMC (see Figure 2d). We were unable to find comparative data in the literature on the effect of pH and monovalent and divalent ion concentrations on the interaction of SMC orthologs with DNA, although for some homologs it was shown that salt inhibits the interaction.

Nevertheless, the data obtained here are important for understanding the functional activity of this protein in living *U. parvum* cells. Taking into account physiological values of these parameters in bacterial cells (pH about 7, about 100 mM KCl and 10 mM MgCl_2_, it can be assumed that SMC/DNA complexes should be formed in *U. parvum* cells.

### DNA-Stimulated ATPase Activity of SMC *U. parvum*

SMC *U. parvum* exhibits ATPase activity stimulated by DNA. Spectrophotometric analysis of the ATPase activity of the *U. parvum* SMC protein using the malachite green assay confirmed the assumption that this protein hydrolyzes ATP in the presence of DNA. Based on a linear approximation of the relationship corresponding to standards with known values of free phosphate concentration, the rate of ATP hydrolysis by the SMC *U. parvum* protein was calculated. It was determined that one molecule of SMC protein (monomer) hydrolyzes 3.8 ± 1.1 (mean ± standard deviation) ATP molecules per minute. In addition, experiments showed that the SMC protein of *U. parvum* is also able to hydrolyze ATP in the absence of DNA, but the rate of hydrolysis is about 2 times lower. These calculations were made assuming that hydrolysis of one molecule of ATP releases one molecule of free phosphate.

Thus, as previously for the SMC protein of *B. subtilis*, we were able to demonstrate DNA-stimulated ATPase activity in this work. At the same time, the ATPase activity of SMC *U. parvum* is about an order of magnitude lower than that of SMC *B. subtilis* (about 30 molecules per minute) [28] and comparable to that of MukB [23].

### DNA Binding of *U. parvum* SMC Prevents DNA Straightening by Molecular Combing

The interaction of *U. parvum* SMC with DNA prevents the latter from straightening by molecular combing. Using the molecular combing method, it was demonstrated that the *U. parvum* SMC protein binds to DNA, preventing it from straightening on the coverslip under the influence of the surface tension force acting on DNA in the liquid-air phase transition zone. It can be seen that DNA straightening is not observed both in the absence of ATP and in the presence of 1 mM ATP (see Figure 3). This result is consistent with the results obtained by electrophoretic mobility shift assay, as it confirms the binding of SMC *U. parvum* c DNA. In addition, the molecular combing method allows us to conclude that this protein compacts one or more DNA molecules, apparently by forming loops from one molecule or by cross-linking several DNA molecules simultaneously.

**Figure 3:**
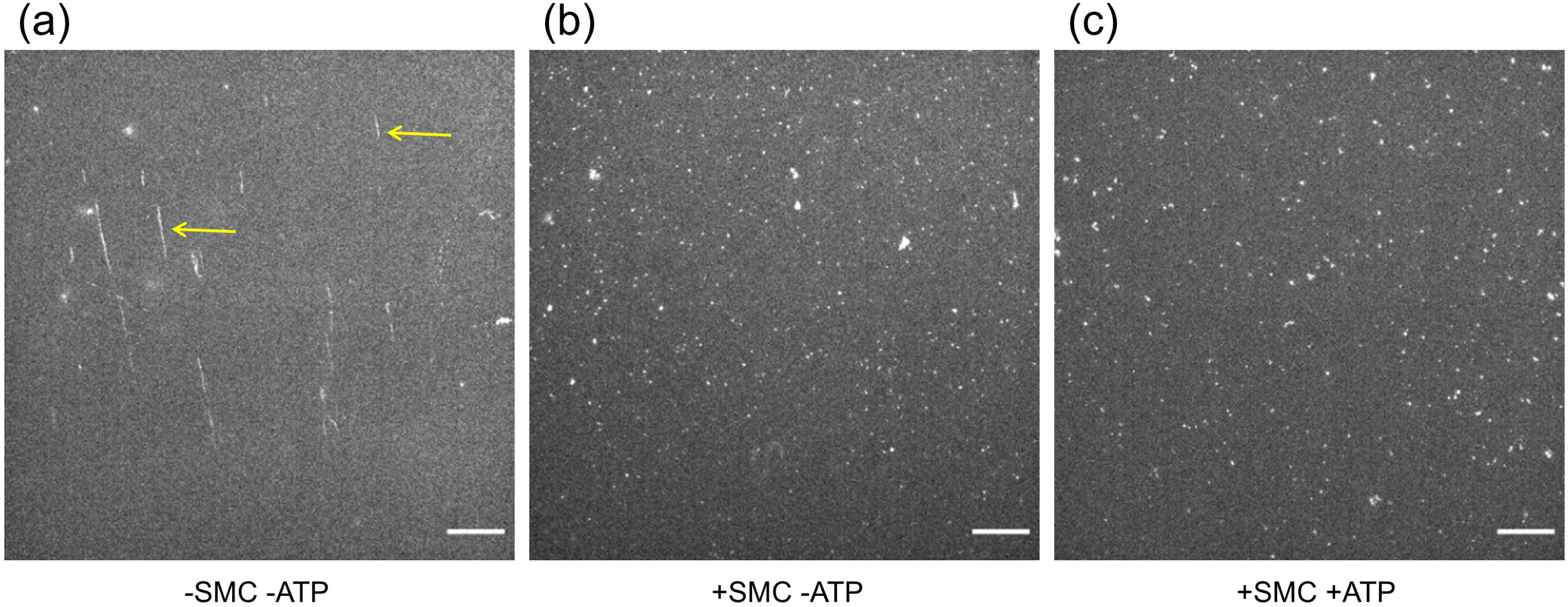
Visualization of λ bacteriophage DNA stretched by molecular combing and stained with fluorescent dye YOYO-1 using fluorescence microscopy. (a) λ DNA incubated without *U. parvum* SMC, showing straightened DNA molecules indicated by arrows. (b) λ DNA incubated with *U. parvum* SMC in the absence of ATP. (c) λ DNA incubated with *U. parvum* SMC and 1 mM ATP. Scale bar: 15 μm

### DNA Shortening by *U. parvum* SMC in Single-Molecule Experiments

*U. parvum* SMC shortens DNA molecules under single-molecule experimental conditions. Using laser tweezers, it was demonstrated that the *U. parvum* SMC protein binds to double-stranded DNA molecules attached at the ends to microspheres, as incubation of a relaxed (unstretched) DNA molecule with SMC without the addition of ATP leads to subsequent DNA shortening at the same tension force compared to the original DNA molecule (see Figure 4). It is interesting to note that incubation of the stretched DNA molecule with SMC solution does not result in a similar effect. Based on this observation, we can make an assumption that in the case of relaxed DNA molecule, the SMC is able to form spatial complexes with it (possibly, to form loops). Probably, SMC forms cross-links between different parts of DNA, which leads to shortening of the latter. This observation is consistent with other results of this work and indicates high stability of SMC/DNA complexes. It is interesting to note that DNA shortening occurred in the absence of ATP, which is similar to the result obtained by molecular combing (see Figure 3).

**Figure 4:**
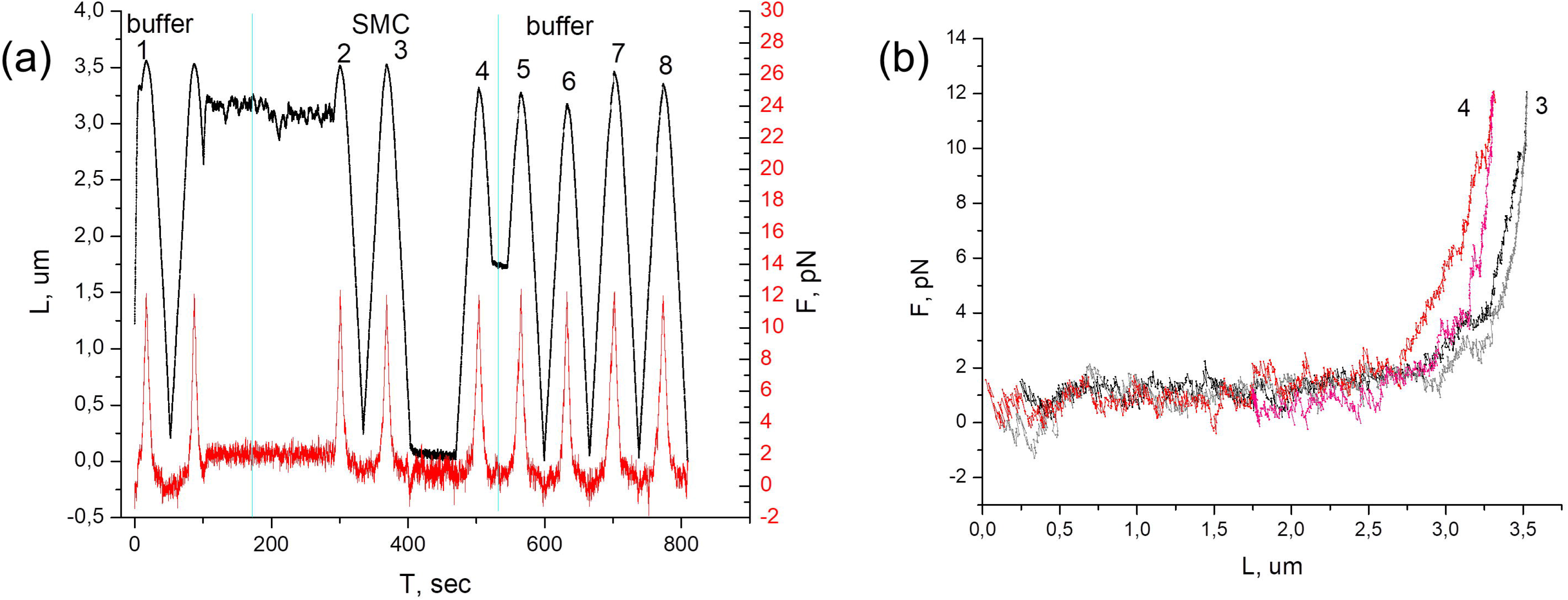
Visualization of *U. parvum* SMC interaction with DNA under single-molecule experimental conditions using laser tweezers. A linear DNA molecule with a length of 11070 bp (prlC) was utilized as substrate in a solution containing Tris-HCl buffer (pH=7.5, 25 mM), 7.5 mM KCl, 2.5 mM MgCl_2_, and 0.02% BSA. (a) An example of a single-molecule experiment showing a DNA molecule subjected to stretching and relaxation upon incubation with SMC. The time dependence of the molecule length (black curve, scale on the left) and its tension force (red curve, scale on the right) are depicted. Blue vertical lines denote the time intervals during which the DNA molecule was exposed to a solution without SMC (buffer) and with SMC protein (SMC). Numbers indicate successive acts of DNA elongation and relaxation. (b) Examples of tension force versus DNA length (force-extension curves) based on the dependencies shown in (a). Curve (3) corresponds to the act of elongation (3) of the DNA molecule after its incubation in the stretched state in the presence of SMC in panel (a), while curve (4) corresponds to the same after incubation in the relaxed state. It is observed that in the latter case, the DNA molecule became shorter at the same tension force

In this work, we were unable to demonstrate active extrusion of loops by the SMC protein of *U. parvum* using laser tweezers and using another single-molecule method, namely the analysis of U-shaped DNA molecules in the liquid flow (data not shown). The most likely reason for this is our use of only the SMC ATPase rather than the whole SMC-ScpAB complex. Probably, in the absence of kleisin (ScpA) and KITE (ScpB), SMC is able to bind to DNA and ATP, hydrolyze ATP; ATP binding and/or hydrolysis somehow stimulate the formation of DNA spatial structures (i.e. SMC cross-links between different segments of one or more DNA molecules), which follows from the data obtained by EMSA (see Figure 1a). Interestingly, the ability to compact genomic DNA independently of ATP binding and hydrolysis was recently demonstrated for the SMC protein of *B. subtilis* under *in vivo* conditions [29], ScpAB modulate this compactization. A similar property was shown for MukB protein - under *in vitro* conditions this protein compacted DNA in the absence of ATP [12]. Given the relatively low ATP hydrolysis rate by the *U. parvum* SMC protein, it is important to explore the connection between its ATPase activity and its ability to compact DNA in the future.

## Conclusions

1. *U. parvum* SMC binds to double-stranded circular and linear DNA.
2. The interaction of *U. parvum* SMC with DNA is inhibited at high KCl, MgCl_2_ concentrations (above 100 mM), as well as at high pH values (above 7).
3. The interaction of *U. parvum* SMC with DNA rises with increasing ATP concentration.
4. *U. parvum* SMC demonstrates DNA-dependent ATPase activity. One SMC monomer hydrolyzes 3.8 ATP molecules per minute.
5. *U. parvum* SMC shortens the DNA as evidenced by molecular combing and laser tweezers data.

## Acknowledgments

The research was supported by the Russian Science Foundation, project No. 22-74-00072 to Morozova N.E. The work was carried out using scientific equipment of the Center of Shared Usage “The Analytical Center of Nano- and Biotechnologies of SPbPU”. The authors are grateful to G.Y. Fisunov for providing the U. parvum strain and I.E. Vishnyakov for his thoughtful assistance in the realization of this work.

## Author Contributions

N.R.: Investigation, Formal analysis, Writing; V.S.: Investigation, Writing; D.B.: Investigation, Formal analysis; A.A.: Investigation, Formal analysis; N.M.: Funding acquisition; A.V.: Supervision, Writing.

